# Nuclear and mitochondrial genomes of the hybrid fungal plant pathogen *Verticillium longisporum* display a mosaic structure

**DOI:** 10.1101/249565

**Authors:** Jasper R.L. Depotter, Fabian van Beveren, Grardy C.M. van den Berg, Thomas A. Wood, Bart P.H.J. Thomma, Michael F. Seidl

**Affiliations:** Laboratory of Phytopathology, Wageningen University, Droevendaalsesteeg 1, 6708 PB Wageningen, The Netherlands; Department of Crops and Agronomy, National Institute of Agricultural Botany, Huntingdon Road, CB3 0LE Cambridge, United Kingdom

**Keywords:** whole-genome duplication, allopolyploidy, genomic rearrangement, Verticillium stem striping, Verticillium wilt, *Brassica*

## Abstract

Allopolyploidization, genome duplication through interspecific hybridization, is an important evolutionary mechanism that can enable organisms to adapt to environmental changes or stresses. This increased adaptive potential of allopolyploids can be particularly relevant for plant pathogens in their quest for host immune response evasion. Allodiploidization likely caused the shift in host range of the fungal pathogen plant *Verticillium longisporum*, as *V. longisporum* mainly infects Brassicaceae plants in contrast to haploid *Verticillium* spp. In this study, we investigated the allodiploid genome structure of *V. longisporum* and its evolution in the hybridization aftermath. The nuclear genome of *V. longisporum* displays a mosaic structure, as numerous contigs consists of sections of both parental origins. *V. longisporum* encountered extensive genome rearrangements, whereas the contribution of gene conversion is negligible. Thus, the mosaic genome structure mainly resulted from genomic rearrangements between parental chromosome sets. Furthermore, a mosaic structure was also found in the mitochondrial genome, demonstrating its bi-parental inheritance. In conclusion, the nuclear and mitochondrial genomes of *V. longisporum* parents interacted dynamically in the hybridization aftermath. Conceivably, novel combinations of DNA sequence of different parental origin facilitated genome stability after hybridization and consecutive niche adaptation of *V. longisporum*.

## INTRODUCTION

Whole-genome duplication (WGD) is an important evolutionary mechanism that facilitates environmental adaptation (Van de Peer et al. 2017; Mallet 2005). The duplication of genomic content increases genomic plasticity, leading to an augmented adaptive potential of organisms that underwent WGD. Consequently, polyploids have been associated with increased invasiveness (te Beest et al. 2012) and resistance to environmental stresses (Lohaus & Van de Peer 2016). For instance, numerous plant species that survived the Cretaceous-Palaeogene mass extinction, 66 million years ago, underwent a WGD which is thought to have contributed to their increased survival rates (Vanneste et al. 2014a; Vanneste et al. 2014b). Both genome copies involved in WGD may have the same species origin, i.e. autopolyploidization, or originate from different species as a result of interspecific hybridization, i.e. allopolyploidization. In general, allopolyploids are believed to have a higher adaptive potential than autopolyploids due to the increased genetic divergence between the chromosome sets.

The impact of allopolyploidization has mainly been investigated in plants, as approximately a tenth of all plant species consists of allopolyploids (Barker et al. 2015). In contrast, allopolyploidization in fungi is far less intensively investigated (Campbell et al. 2016). Nonetheless, allopolyploidization impacted the evolution of numerous fungal species, including the economically important baker’s yeast *Saccharomyces cerevisiae* (Marcet-Houben & Gabaldón 2015). The increased adaptive potential enabled allopolyploid fungi to develop desirable traits that can be exploited in industrial bioprocessing (Peris et al. 2017). For instance, at least two recent hybridization events between *S. cerevisiae* and its close relative *Saccharomyces eubayanus* gave rise to *Saccharomyces pastorianus*, a species with high cold tolerance and good maltose/maltotriose utilization capabilities, which is exploited in the production of lager beer that requires barley to be malted at low temperatures (Gibson & Liti 2015).

Allopolyploid genomes experience a so-called “genome shock” upon hybridization, inciting major genomic reorganizations that can manifest by genome rearrangements, extensive gene loss, transposon activation, and alterations in gene expression (Doyle et al. 2008). These early stage alterations are primordial for hybrid survival, as divergent evolution is principally associated with incompatibilities between the parental genomes (Matute et al. 2010). Allopolyploids benefit from a thorough re-organization where negative interactions between the parental genomes are purged. Frequently, heterozygosity is lost for many regions in the allopolyploid genome (Mixão & Gabaldón 2018). This can be a result of the direct loss of a duplicated gene copy through deletion or gene conversion, a process where one of the copies substitutes its homeologous counterpart (McGrath et al. 2014). Gene conversion and the homogenization of complete chromosomes played a pivotal role in the evolution of the osmotolerant yeast species *Pichia sorbitophila* (Louis et al. 2012). In total, two of its seven chromosome pairs consist of partly heterozygous, partly homozygous sections, whereas two chromosome pairs are completely homozygous. Gene conversion may eventually result in chromosomes consisting of sections of both parental origins as “mosaic genomes” (Stukenbrock et al. 2012). However, mosaic genomes can also arise through recombination between chromosomes of the different parents, such as in the hybrid yeast *Zygosaccharomyces parabailii* (Ortiz-Merino et al. 2017).

Plant pathogens are often thought to evolve while being engaged in arms races with their hosts; pathogens evolve to evade host immunity while plant hosts attempt to intercept pathogen ingress (Cook et al. 2015). Due to the increased adaptation potential, allopolyploidization has been proposed as a potent driver in pathogen evolution (Depotter et al. 2016b). Allopolyploids often have different pathogenic traits than their parental lineages, such as higher virulence (Husson et al. 2015; Brasier & Kirk 2001) and shifted host ranges (Inderbitzin et al. 2011b; Zeise & Tiedemann 2002). Within the fungal genus *Verticillium*, allodiploidization resulted in the emergence of a novel pathogen on brassicaceous plants; *Verticillium longisporum* (Inderbitzin et al. 2011b; Depotter et al. 2017b). Similar to haploid *Verticillium* spp., *V. longisporum* is thought to have a predominant asexual reproduction as a sexual cycle has never been described and populations are not outcrossing (Depotter et al. 2017b; Short et al. 2014). *V. longisporum* is sub-divided into three lineages, each representing a separate hybridization event (Inderbitzin et al. 2011b). The economically most important lineage is A1/D1 that originates from hybridization between *Verticillium* species A1 and D1 that have hitherto not been found in their haploid states. *V. longisporum* lineage A1/D1 is the main causal agent of Verticillium stem striping on oilseed rape (Novakazi et al. 2015) and its economic importance as emerging pathogen is increasing worldwide (Depotter et al. 2017a). A recent study revealed that lineage A1/D1 can be further divided into two genetically distinct populations, which have been named ‘A1/D1 West’ and ‘A1/D1 East’ after their geographic occurrence in Europe (Depotter et al. 2017b). Nevertheless, both populations were shown to originate the same hybridization event (Depotter et al. 2017b).

*V. longisporum* is assumed to have largely conserved its allodiploid state as the sizes of its sub-genomes resemble those of haploid *Verticillium* spp. (Depotter et al. 2017b; Shi-Kunne et al. forthcoming; Fogelqvist et al. 2018). Nevertheless, not all genes are present in heterozygous copies, as its nuclear ribosomal internal transcribed spacer region is derived only from one of the parents (Inderbitzin et al. 2011b). Here, we investigated the evolution of the allodiploid genome of *V. longisporum* and determined to what extent heterozygosity is lost.

## MATERIAL AND METHODS

### Genome analysis

Genome assemblies of the two *V. longisporum* strains (VLB2 and VL20) and *V. dahliae* strain JR2 were previously published (Faino et al. 2015; Depotter et al. 2017b). Telomeric regions were determined based on the telomeric repeat pattern: TAACCC/GGGTTA (minimum three repetitions). Furthermore, additional repeats were identified and characterized using RepeatModeler (v1.0.8). De-novo-identified repeats were combined with the repeat library from RepBase (release 20170127) (Bao et al. 2015). The exact coordinates of the repeats were extracted with the software RepeatMasker (v4.0.6) (Smit et al. 2015). Genome-wide sequence identities between *Verticillium* strains were calculated with dnadiff (Kurtz et al. 2004). Homologous genes were retrieved by nucleotide BLAST (v2.2.31+). Here, only hits with a minimal coverage of 80% with each other were selected.

### Gene annotation

*Verticillium* isolates JR2, VLB2 and VL20 were grown for 3 days in potato dextrose broth. Total RNA was extracted based on TRIzol RNA extraction (Simms et al. 1993). cDNA synthesis, library preparation (TreSep RNA-Seq short-inser library), and Illumina sequencing (single-end 50bp) was performed at the Beijing Genome Institute (BGI, Hong Kong, China). In total, ~2Gb of filtered reads were mapped on the *Verticillium* genomes using TopHat (Trapnell et al. 2009) with Bowtie2 (Langmead & Salzberg 2012). Gene annotation was performed with the BRAKER1 1.9 pipeline (Hoff et al. 2016) using GeneMark-ET (Lomsadze et al. 2014) and AUGUSTUS (Stanke et al. 2008). Predicted genes with internal stop codons were removed from the analysis.

### Parental origin determination

Sub-genomes were divided based on the differences in sequence identities between species A1 and D1 with *V. dahliae*. *V. longisporum* genomes of VLB2 and VL20 were aligned to the complete genome of *V. dahliae* JR2 using NUCmer from the MUMmer package v3.23 (Kurtz et al. 2004; Faino et al. 2015). Here, only 1-to-1 alignments longer than 10kb and a minimum of 80% identity were retained. Subsequent alignments were clustered together. The average nucleotide identity was determined for every cluster and used for sub-genome segregation.

The parental origin determination based on sequence identities of the exonic regions of genes, which was performed by BLAST (v2.6.0+). Here, hits with a minimum subject and query coverage of 80% were used. Furthermore, similar to Louis et al. (2012), differences in GC-content between homolog genes present in two copies were calculated accordingly:

dGCgene = 2*GCgene / (GCgene + GChomolog)

GCgene = GC% of gene

GChomolog = GC% of homolog

dGCgene = GC ratio from the mean GC% value

### Gene conversion and genomic rearrangements

Double copy genes were retrieved by nucleotide BLAST (v2.6.0+) and the sequence identity determined. Here, hits with a minimum subject and query coverage of 80% were used. Nucleotide BLAST (v2.6.0+) was also used to find the corresponding homologous genes in *V. longisporum* strains VLB2 and VL20, which were present in two copies in both strains.

The VLB2 genome assembly was aligned to VL20 to find breaks in synteny using NUCmer from the MUMmer package v3.23 (Kurtz et al. 2004). Subsequent alignments were clustered if they aligned to the same contig with the same orientation and order as the reference genome. In order to confirm the breaks in synteny, filtered *V. longisporum* reads of VLB2 were aligned to the *V. longisporum* VL20 genome with the Burrows-Wheeler Aligner (BWA) and further processed with the samtools package (v1.3.1) (Li et al. 2009). Breaks in syntenic clusters were then visualized and determined using the R package Sushi (Phanstiel et al. 2014) and the Integrative Genomics Viewer (Robinson et al. 2011). The association between breaks with repeats was tested through permutation. First, the fraction of breaks flanked by repeats was determined. Here, breaks were assigned to reside in a “repeat-rich” region if a 1 kb window around the break consisted for more than 10% of repeats. Then, the *V. longisporum* VL20 genomes was divided into windows of 1 kb using BEDTools (v2.26.0) to calculate the significance of the break/repeat association (Quinlan & Hall 2010). In total, 10,000 permutations were executed with the same amount of windows to determine the random distribution of repeat-rich regions.

### Phylogenetic tree construction

The phylogenetic tree of the nuclear DNA was based on the nucleotide sequences of the ascomycete set BUSCO orthologs present in clade Flavnonexudans *Verticillium* spp. (Simão et al. 2015). In total, 1,194 genes were included and concatenated for tree construction. The mitochondrial phylogenetic tree was based on the nucleotide sequence of complete or partial mitochondrial genomes. Nuclear and mitochondrial genomes of *Verticillium* spp. other than *V. longisporum* were previously sequenced and assembled (Shi-Kunne et al. forthcoming; Faino et al. 2015; Jelen et al. 2016). Whole-genome alignments for tree construction were performed by mafft (v7.271) (default settings) (Katoh & Standley 2013; Katoh et al. 2002), and subsequently the likelihood phylogenetic tree was reconstricted using RAxML with the GTRGAMMA substitution model (v8.2.0) (Stamatakis 2014). The robustness of the inferred phylogeny was assessed by 100 rapid bootstrap approximations.

## RESULTS

### *V. longisporum* displays a mosaic genome structure

The genomes of two *V. longisporum* strains were analysed to investigate the impact of hybridization on the genome structure. Previously, *V. longisporum* strains VLB2 and VL20, belonging to ‘A1/D1 West’ and ‘A1/D1 East’, were sequenced with the PacBio RSII platform and annotated into genomes of 72.9 and 72.3 Mb in size, respectively (Depotter et al. 2017b). These genome sizes exceed double the amount of the telomere-to-telomere sequenced *V. dahliae* strains JR2 (36.2 Mb) and VdLs17 (36.0 Mb) (Faino et al. 2015). Despite the previous observation that the two *V. longisporum* strains belong to distinct populations within the A1/D1 lineage, the genomes show an extremely high degree of sequence identity (99.91%). We used RepeatModeler (V1.0.8) in combination with RepeatMasker to determined that 14.28 and 13.90% of the *V. longisporum* strain VLB2 and VL20 genomes is composed of repeats, respectively (Table 1) (Smit et al. 2015). Intriguingly, this is more than double the repeat content as in *V. dahliae* strain JR2, for which 6.49% of the genome was annotated as repeat using the same methodology. The *V. longisporum* genomes were also screened for telomere-specific repeats (TAACCC/GGGTTA) to estimate the number of chromosomes. In total, 29 and 30 telomeric regions were found in the VLB2 and VL20 genomes, respectively, that were consistently situated at the end of contigs, suggesting that *V.longisporum* contains at least 15 chromosomes (Table 1). Six out of 45 and four out of 44 contigs in strains VLB2 and VL20, respectively, were flanked on both ends by telomeric repeats and therefore likely represent complete chromosomes (Table 1). For comparison, *V. dahliae* strains contain 8 chromosomes (Faino et al. 2015).

**Table 1:** Comparison *V. longisporum* and *V. dahliae* genomes.

|  | *V. longisporum* | *V. longisporum* | *V. dahliae* |
| --- | --- | --- | --- |
|  | **VLB2** | **VL20** | **JR2** |
| **Genome size** | 72.9 Mb | 72.3 Mb | 36.2 Mb |
| **Number of contigs** | 45 | 44 | 8 |
| **Complete chromosomes** | 6 | 4 | 8 |
| **Number of genes** | 19,123 | 18,784 | 9,909 |
| **Telomere regions** | 29 | 30 | 16 |
| **Repeat content** | 14.28 | 13.90 | 6.49 |
| **BUSCO completeness** | 99.0% | 98.3% | 98.7% |

In allopolyploid organisms, parental origin determination is elementary to investigate genome evolution in the hybridization aftermath. As species D1 is phylogenetically closer related, and consequently has a higher sequence identity, to *V. dahliae* than species A1, *V. longisporum* genomic regions were previously provisionally assigned to either species D1 or A1 (Depotter et al. 2017b). Here, we determined the parental origin of *V. longisporum* genomic regions more precisely. The difference in phylogenetic distance of species A1 and D1 to *V. dahliae* caused that *V. longisporum* genome alignments to *V. dahliae* displayed a bimodal distribution with one peak at 93.1% and another peak at 98.4% sequence identity that represent the two parents with a dip in between at 96.0% (Figure S1). In order to separate the two sub-genomes, regions with an average sequence identity to *V. dahliae* <96% were assigned to species A1, whereas regions with an identity of ≥96% were assigned to species D1 (Figure 1). In this manner, 36.2 Mb of *V. longisporum* strain VLB2 was assigned to species A1 and 35.7 Mb to species D1. For *V. longisporum* strain VL20, 36.3 Mb was assigned to species A1 and 35.2 Mb to species D1. Only 1.0 and 0.8 Mb of strains VLB2 and VL20 could not be aligned to *V. dahliae* and thus remained unassigned, respectively.

**Fig. 1:**
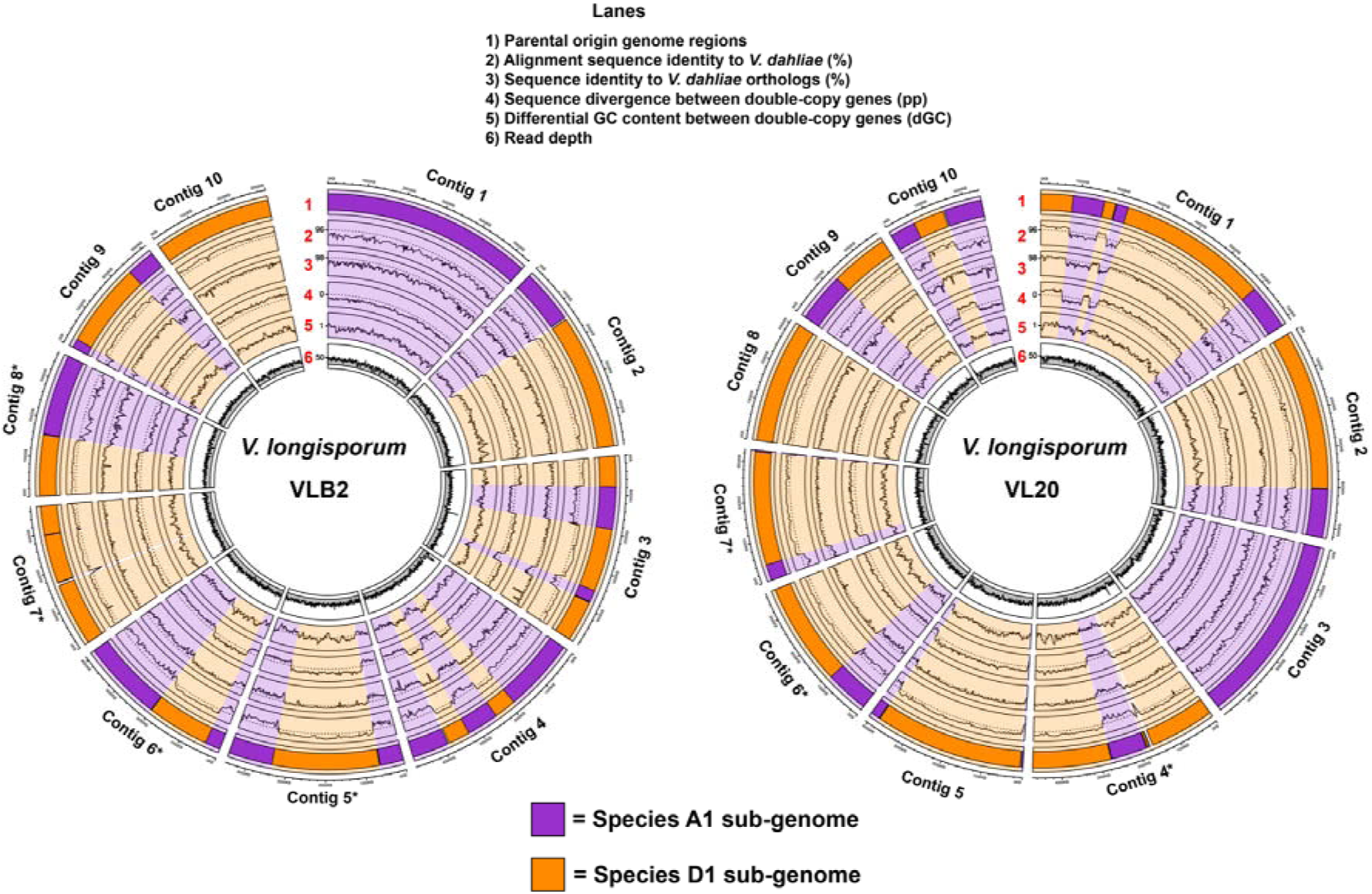
Determination of parental origin of *Verticillium longisporum* genome sections. The ten largest contigs of the genome assemblies of *V. longisporum* strains VLB2 and VL20 are depicted. Lane 1: regions with a sequence identity of at least 96% were assigned to Species D1 (orange), whereas ones with lower sequence identity to Species A1 (purple). Lane 2: sequence identity of *V. longisporum* alignments to *V. dahliae*. Lane 3: Sequence identity between exonic regions of *V. longisporum* and *V. dahliae* orthologs. Lane 4: Difference in sequence identity in percent point (pp) between exonic regions of *V. longisporum* double-copy genes. Only gene pairs with an ortholog in *V. dahliae* are depicted. Alleles with a higher identity to *V. dahliae* are depicted as a positive pp difference, whereas the corresponding homolog as a negative pp difference. Lane 5: the relative difference in GC content (dGC) between genes in double copy. Lane 6: Read depth with non-overlapping windows of 10 kb. Data points of lanes 3-5 represent the average value of a window of eleven genes, which proceed with a step of one gene.

To trace the chromosome sets of the original parents of the hybrid, the parental origin of individual contigs was determined. In total, 8 of the 10 largest contigs of *V. longisporum* strain VLB2 as well as strain VL20 consist of regions originating from both species A1 and species D1 (Figure 1). Thus, parental chromosome sets cannot be separated from one another as *V. longisporum* apparently evolved a mosaic genome structure in the hybridization aftermath.

### Genomic rearrangements are responsible for the mosaic genome

Typically, a mosaic structure of a hybrid genome can originate from gene conversion or from chromosomal rearrangements between DNA strands of different parental origin (Mixão & Gabaldón 2018). To analyse the extent of gene conversion, genes were predicted for the *V. longisporum* strains VLB2 and VL20. To aid gene annotation with the BRAKER1 1.9 pipeline (Hoff et al. 2016), ~2 Gb of filtered RNA-seq reads were generated from fungal cultures in potato dextrose broth. In total, 19,123 and 18,784 genes were predicted for *V. longisporum* strains VLB2 and VL20 respectively, which is ~90% higher than the amount of genes that were predicted for *V. dahliae* strain JR2 in this manner (9,909 genes) (Table 1). As expected, the divergence of species A1 and D1 was also reflected at the level of gene sequences based on sequence identity and GC-content (Figure 1, S1). In total, 9,531 and 9,402 genes were assigned to the species A1 sub-genome of the strains VLB2 and VL20, respectively, whereas the number of genes in the species D1 sub-genomes was 9,468 and 9,243 for these strains, respectively (Figure 2). Thus, the amount of genes is similar in the two sub-genomes for both *V. longisporum* strains. Over 80% of the *V. longisporum* genes are present in two copies whereas, similar to *V. dahliae*, almost all genes (97-98%) are present in one copy within each of the *V. longisporum* sub-genomes. Moreover, of the 7,620 genes that are present in two copies in VLB2 and VL20, only 5 genes were found to be highly identical (<1%, nucleotide sequence identity) in VLB2, whereas the corresponding gene pair in VL20 was more diverse (>1%, nucleotide sequence identity) (Figure 3). In *V. longisporum* strain VL20, no highly identical copies were found that are more divergent in VLB2. Collectively, these findings indicate that most *V. longisporum* genes have a copy of a different parental origin and that gene conversion played a minor role in during evolution of the mosaic genome.

**Fig. 2:**
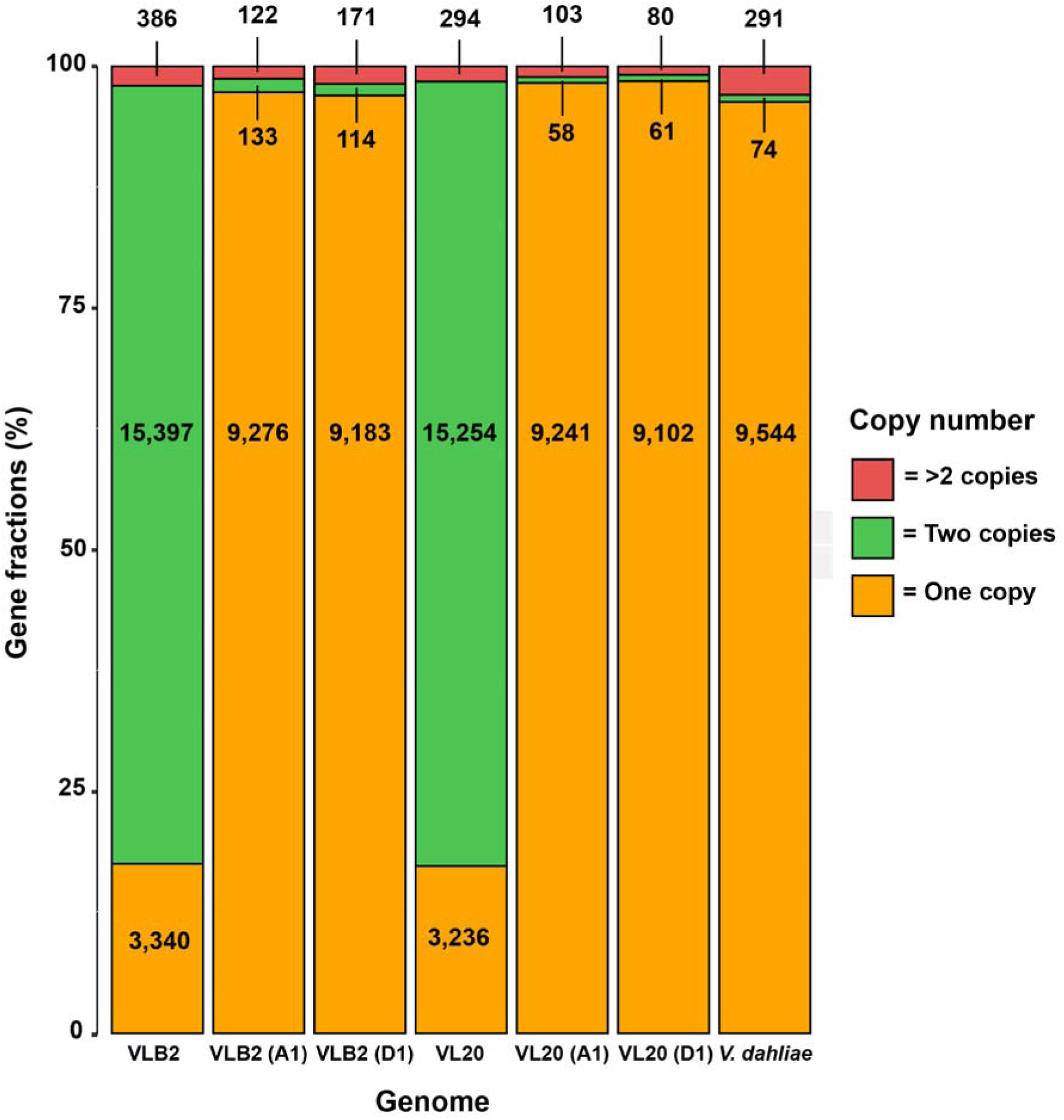
Gene copy number distribution within *Verticillium* (sub-)genomes. “(A1)” and “(D1)” represent species A1 and D1 sub-genomes, respectively, of the *V. longisporum* strains VLB2 and VL20. For *V. dahliae*, the strain JR2 was used.

**Fig. 3:**
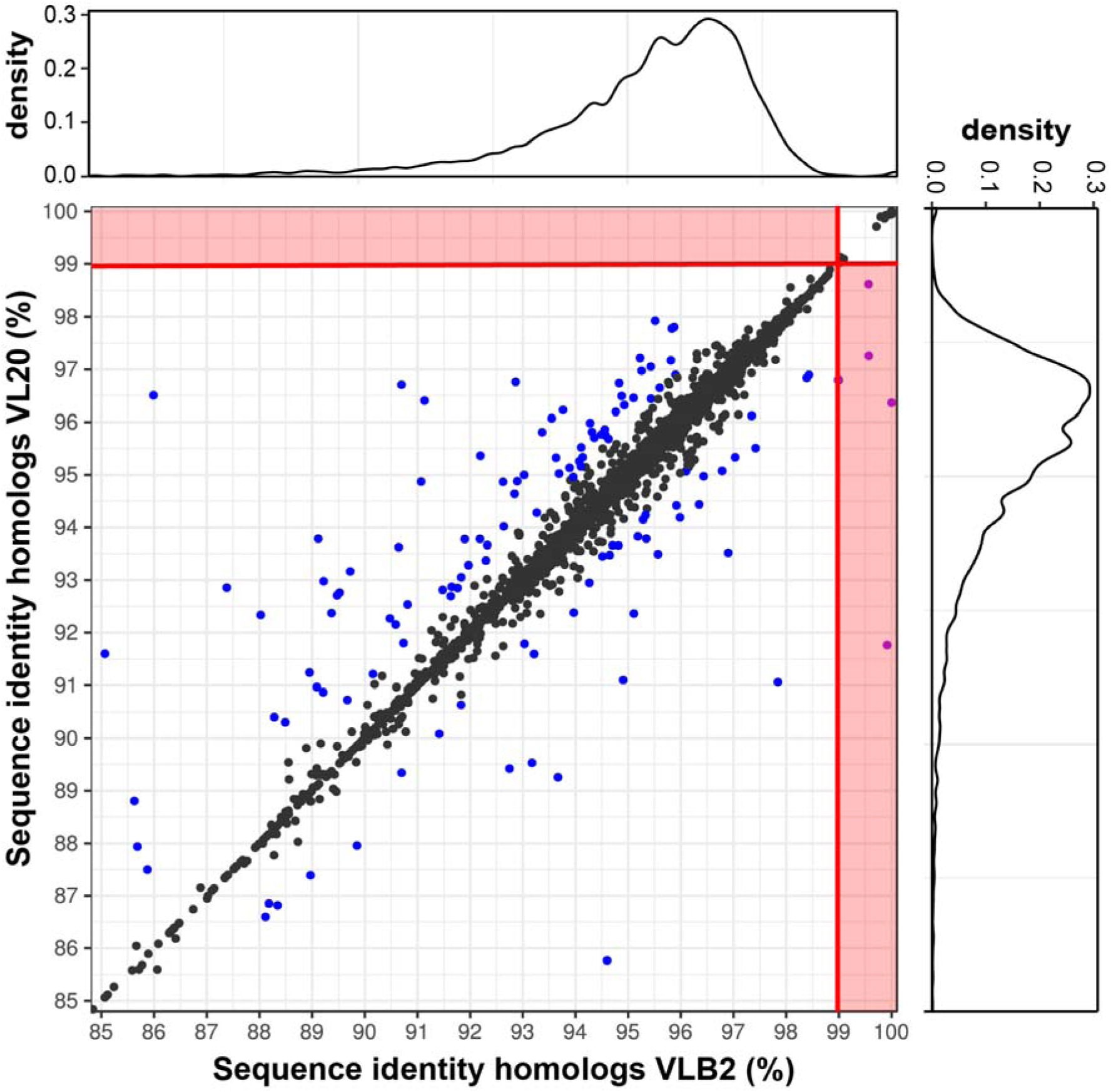
The contribution of gene conversion to *V. longisporum* genome evolution. Sequence identities between double-copy genes, present in *V. longisporum* VLB2 and VL20, are depicted. Gene pairs that encountered gene conversion (purple dots in the red zones) have sequence divergence of more than one percent in one *V. longisporum* strain and less than one percent in the other strain. In other cases, pairs that differ less than one percent are depicted as a black dot, whereas a difference higher than one percent is depicted as a blue dot.

Considering that gene conversion played a minor role during genome evolution, the mosaic genome structure of *V. longisporum* is likely to originate from rearrangements between the chromosomes of different parental origin. To identify the location of genomic rearrangements, the genome of *V. longisporum* strain VLB2 was aligned to that of strain VL20 (Figure 4). Extensive rearrangements were observed between the two *V. longisporum* strains, as 87 putative syntenic breaks were found. In order to confirm these synteny breaks, individual long-reads of VLB2 were aligned to the VL20 genome assembly to assess if breaks were supported by read mapping (Figure S2). In total, 60 synteny breaks could be confirmed by read mapping. As genomic rearrangements are often associated with repeat-rich genome regions, the synteny break points were tested for their occurrence in repeat-rich regions. In total, 34 of the 60 (57%) confirmed synteny break points were flanked by repeats, which is significantly more than what would be expected from random sampling (mean = 18.5%, σ = 0.05%) (Figure S3). In conclusion, it appears that genomic rearrangements, rather than gene conversion, are the main driver behind the mosaic structure of the *V. longisporum* genome.

**Fig. 4:**
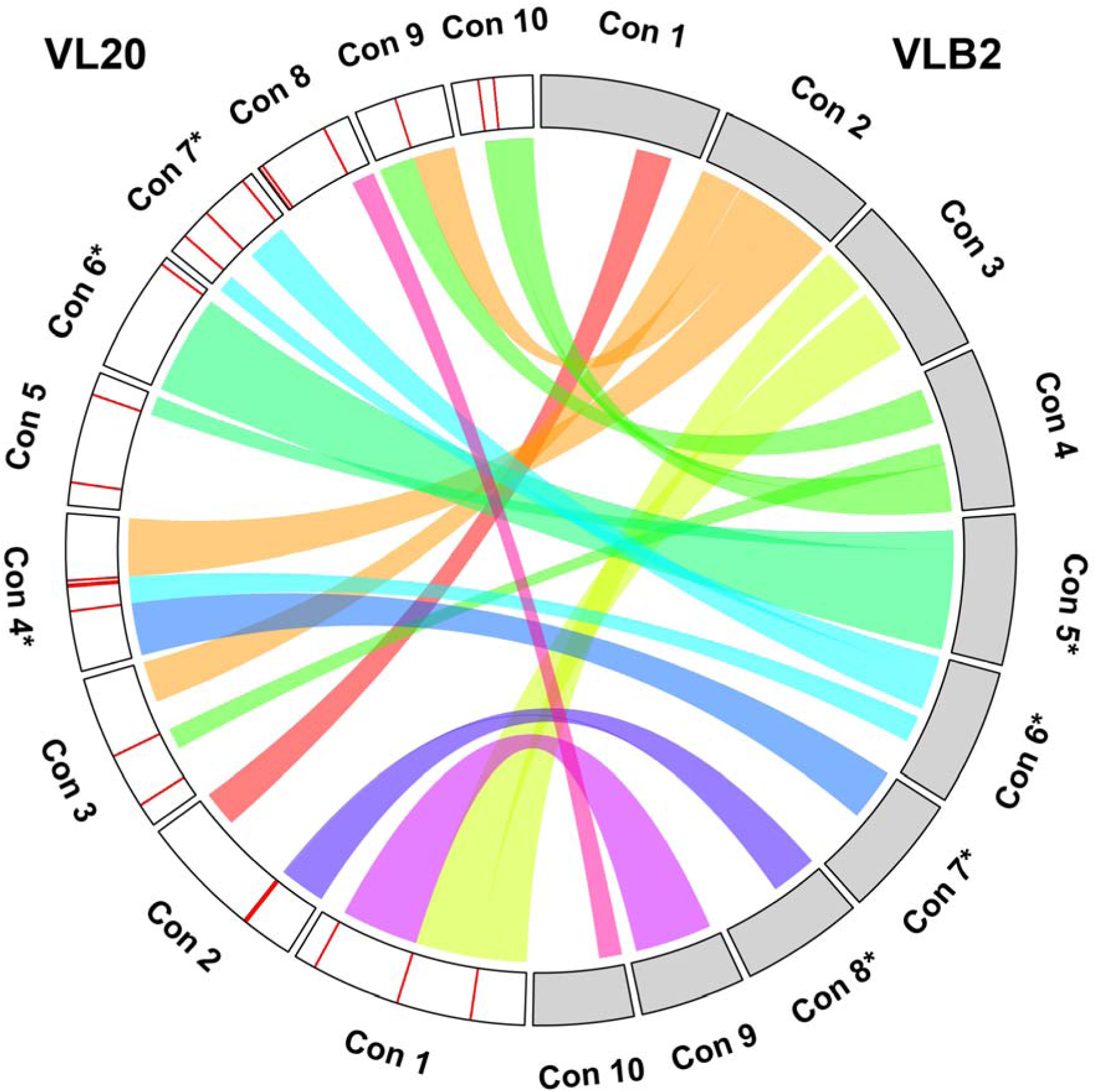
The contribution of genomic rearrangements to *V. longisporum* genome evolution. The ten largest contigs of the *V. longisporum* strains VLB2 (displayed in grey) and VL20 (displayed in white) are depicted with complete chromosomes indicated by asterisks. Ribbons indicate syntenic genome regions between the two strains. Red bars on the contigs indicate synteny breaks that are confirmed by discontinuity in alignment of VLB2 reads to VL20 genome.

### *V. longisporum* loses heterozygosity through deletions

In each of the *V. longisporum* isolates, 17% of the genes occur only in a single copy. Although gene conversion played a minor role in the hybridization aftermath, loss of heterozygosity may occur through gene loss or, alternatively, single-copy genes may originate from parent-specific contributions. However, as 12% of the singly copy genes in strain VLB2 are present in two copies in strain VL20, and 16% of the single copy genes in VL20 are present in two copies in VLB2, gene deletion seems to be an on-going process in *V. longisporum* evolution since both strains are derived from the same hybridization event (Depotter et al. 2017b). Thus, in the general absence of gene conversion in the *V. longisporum* genome, loss of heterozygosity is mainly caused by deletions.

### Also the mitochondrial genome has a bi-parental origin

To determine how *Verticillium* species A1 and D1 relate to other species in the clade Flavnonexudans, a phylogenetic tree was constructed based on 1,194 Ascomycota Benchmarking Universal Single-Copy Orthologs (BUSCOs) that are present in all members of *Verticillium* clade Flavnonexudans (Figure 5). Species A1 diverged before the last common ancestor of *V. alfalfa, V. dahliae* and *V. nonalfalfae*. In contrast, species D1 only recently diverged from *V. dahliae* after the last common ancestor of *V. alfalfae*, *V. dahliae* and *V. nonalfalfae*. In addition to genomic DNA, the *V. longisporum* clade Flavnonexudans phylogeny was also determined based on mitochondrial DNA (mtDNA) (Figure 5). The mitochondrial genomes of the haploid clade Flavnonexudans spp. were previously sequenced and found to be between 25-27 kb in size (Shi-Kunne et al. forthcoming; Jelen et al. 2016). The *V. longisporum* mtDNA was assembled along with the nuclear genome of VLB2 and VL20, which resulted in a mitochondrial genome of 26.1 kb. Unanticipatedly, the phylogenetic position of *V. longisporum* based on mtDNA did not correspond to the previously determined phylogenetic positions of either of the parents based on the nuclear DNA (Figure 5), as it appeared that *V. longisporum* diverged more recently from the *V. alfalfae*/*V. nonalfalfae* cluster than *V. dahliae*. As hybrid genome structures may impede truthful phylogenetic resolution (Linder & Rieseberg 2004), we hypothesized that a hybrid origin of the *V. longisporum* mitochondrial genome caused the discrepancy between the phylogenetic trees based on nuclear and mitochondrial DNA. In order to elucidate the hybrid nature, differences in mitochondrial sequence identities of *V. longisporum* or *V. nonalfalfae* compared with *V. dahliae* were used to determine the parental origin of *V. longisporum* mitochondrial genomic regions (Figure 6). As *V. nonalfalfae* diverged more recently from *V. dahliae* than species A1, yet before the divergence of *V. dahliae* and species D1, mtDNA that originates from species A1 and D1 should have lower and higher sequence identity, respectively, with *V. dahliae* than with *V. nonalfalfae*. Indeed, the *V. longisporum* mitochondrial genome consists of sections with higher and lower identity to *V. dahliae* than to *V. nonalfalfae* confirming the hybrid nature of the *V. longisporum* mitochondrial genome (Figure 6). The hybrid nature of the *V. longisporum* mtDNA was furthermore confirmed with a phylogenetic analysis based on a 3.5 kb region that displays 0.7% higher average sequence identity to *V. dahliae* than to *V. nonalfalfae*. This mtDNA region placed *V. longisporum* in the same phylogenetic position of species D1 with a divergence after the last common ancestor of *V. dahliae*, *V. alfalfae* and *V. nonalfalfae* (Figure 6). In contrast, a *V. longisporum* mtDNA region of the same length that has on average 0.4% lower sequence identity to *V. dahliae* than *V. nonalfalfae* placed *V. longisporum* in the same phylogenetic position as species A1 (Figure 6). In conclusion, in addition to the nuclear genome, also the mtDNA of *V. longisporum* displays a mosaic structure after recombination of the DNA of the two individual parents.

**Fig. 5:**
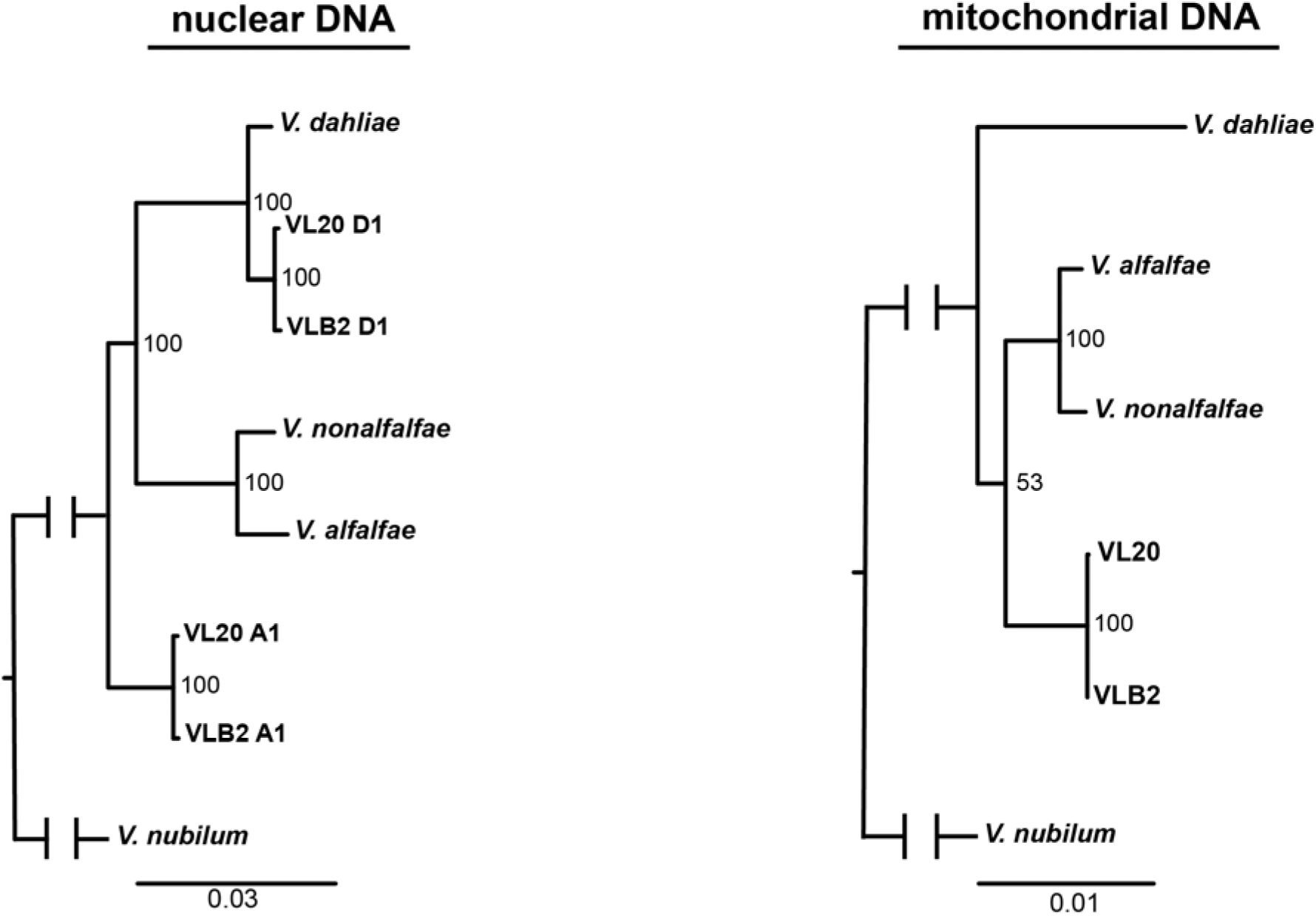
Phylogenetic positioning of *Verticillium longisporum* nuclear sub-genomes and mitochondrial genome to other *Verticillium* species of clade Flavnonexudans. The nuclear phylogenetic tree was constructed with 1,194 orthologous genes, whereas the mitochondrial phylogenetic tree was based on complete mitochondrial genomes. Maximum-likelihood phylogeny analysis of *Verticillium* spp. was rooted on *Verticillium nubilum* and the robustness of the tree was assessed using 100 bootstrap replicates.

**Fig. 6:**
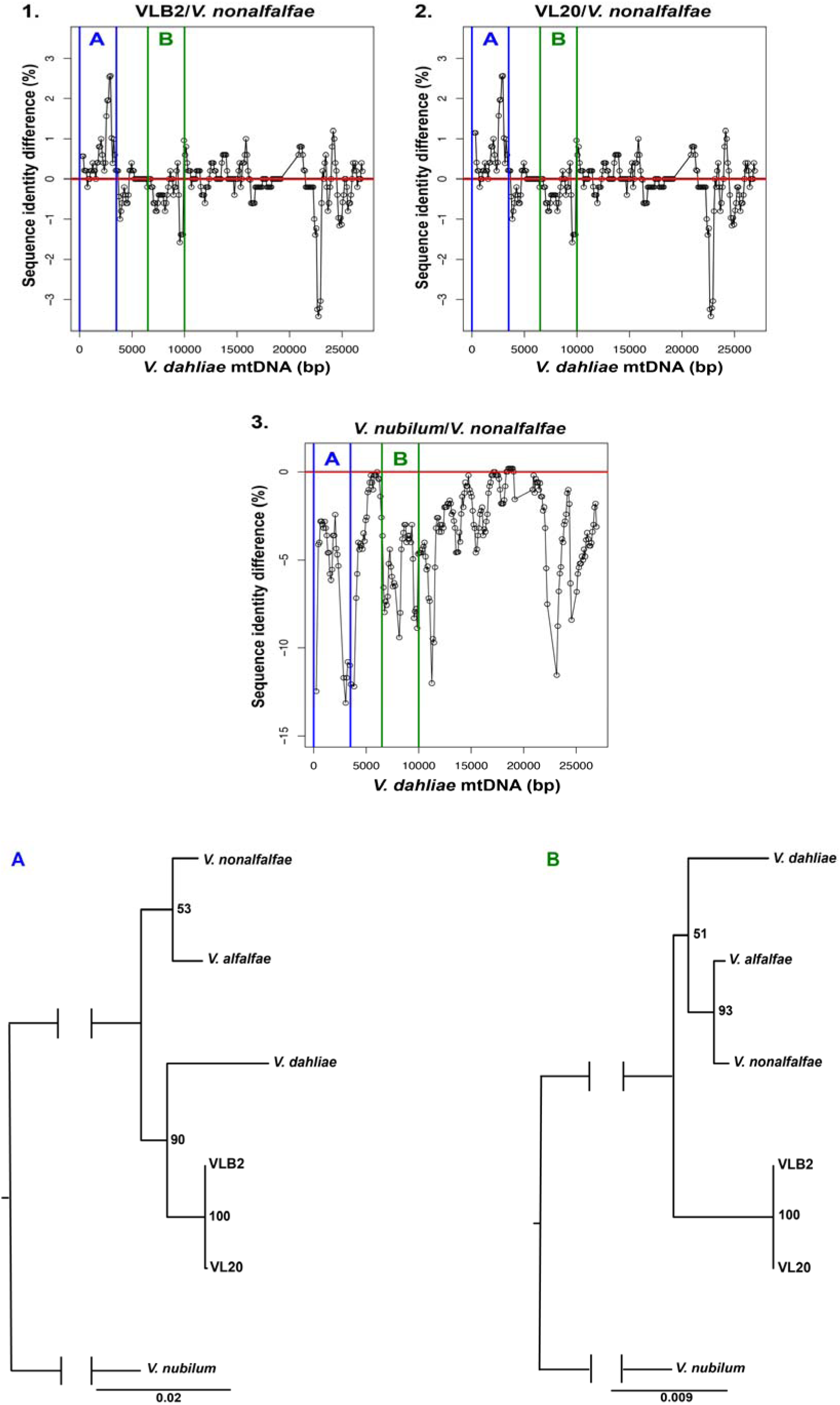
The bi-parental origin to mitochondrial genome of *Verticillium longisporum*. The *V. dahliae* mitochondrial genome was divided in 500 bp sliding windows with 100 bp steps. The differences in sequence identity of these windows with other *Verticillium* genomes were determined for: 1. *V. longisporum* strain VLB2 and *V. nonalfalfae*, 2. *V. longisporum* strain VL20 and *V. nonalfalfae*, and 3. *V. nubilum* and *V. nonalfalfae*. The two phylogenetic trees are constructed with maximum-likelihood based on region A and B of the mitochondrial *V. dahliae* genome, both 3.5 kb in size. The phylogenetic trees were rooted on *Verticillium nubilum* and the robustness of the tree was assessed using 100 bootstrap replicates.

## DISCUSSION

Divergent evolution often fixates genomic incompatibilities between populations, leading to reproductive isolation and eventually even speciation (Seehausen et al. 2014). *Verticillium* species A1 and D1 are distinct with a genome-wide nucleotide divergence of 7.6%, yet both species overcame putative incompatibilities and hybridized into a stable allodiploid (Depotter et al. 2016a). Conceivably, extensive genome alterations occurred after hybridization, facilitating the *V. longisporum* genome to reach a stable equilibrium. The dynamic nature of the *V. longisporum* genome during the hybridization aftermath is displayed by its mosaic structure, not only in the nuclear genome, but even in the mitochondrial genome (Figure 1). Mosaicism in *V. longisporum* is not driven by homogenization that played a negligible role in the hybridization aftermath (Figure 3). Rather, *V. longisporum* mosaic genome structure is caused by extensive genomic rearrangements after hybridization (Figure 4, S2). Genomic rearrangements are major drivers of evolution and facilitate adaptation to novel or changing environments (Seidl & Thomma 2014). This mode of evolution is not specific to the hybrid nature of *V. longisporum* as *V. dahliae* similarly encountered extensive chromosomal reshuffling (de Jonge et al. 2013; Faino et al. 2016). In *V. dahliae*, genomic rearrangements are associated with the occurrence of lineage-specific regions that are derived from segmental duplications, and that are crucial for fungal aggressiveness as they are enriched for transcribed transposable elements and *in planta*-expressed genes (de Jonge et al. 2013; Faino et al. 2016). Genomic rearrangements often result from double-strand DNA breaks that are erroneously repaired based on templates that display high sequence similarity (Seidl & Thomma 2014). As expected, the majority of the synteny breaks between the genomes of *V. longisporum* strains VLB2 and VL20 reside in repeat-rich genome regions (Figure S3) as, due to their abundance, repetitive sequences are more likely to act as a substrate for unfaithful repair (Seidl & Thomma 2014). Nevertheless, 43% of the synteny breaks identified in *V. longisporum* are not associated with repeat-rich regions. However, the presence of two genomes provides orthologous sequences with sufficient identity to mediate unfaithful repair. Double-strand DNA breaks can be generated by transposable elements (TEs). Transposon activity is typically constrained by epigenetic mechanisms such as DNA methylations. However, these constrained may be alleviated upon genome challenges, such as allopolyploidization (Slotkin & Martienssen 2007). The “genome-shock” that *V. longisporum* encountered upon hybridization may have induced TE activity. The higher repeat content of the *V. longisporum* genome compared to the haploid *V. dahliae* may suggest that TE proliferation occurred after hybridization, leading to a modest genome expansion, resulting in a genome size of *V. longisporum* that is more than double when compared with *V. dahliae*, whereas *V. longisporum* contains only 90% more genes (Table 1). Similarly, it has been suggested that hybridization led to TE proliferation in allopolyploid root-knot nematodes, *Meloidogyne* spp., as TEs cover a ±1.7 times higher proportion of their genomes when compared with the closely related, yet non-hybrid, *Meloidogyne hapla* (Blanc-Mathieu et al. 2017).

Whole-genome duplication events are usually followed by extensive gene loss, often leading to reversion to the original ploidy state (Maere et al. 2005). However, the so-called ‘haploidization’ of *V. longisporum* has only proceeded to a limited extent, as 80% of the genes are present in two copies (Figure 2), whereas the haploid *V. dahliae* genome contains only 1% of its genes in two copies. Perhaps *V. longisporum* hybridized only recently, with gene-loss being an on-going process that by now has only progressed marginally, but that will lead to further losses over time. Alternatively, the retention of genes in two copies is of an evolutionary advantage, as the two copies make an additive contribution or their redundancy provides a source for functional divergence (Gu et al. 2002; Blanc & Wolfe 2014). Furthermore, the majority of the two parental genomes may also be retained to maintain genomic balance, as stoichiometric difference of in interacting genes may have detrimental outcomes (Birchler & Veitia 2012).

Hybridization can lead to incongruences between phylogenetic trees based on genome sections of different parental origin (Linder & Rieseberg 2004). High-resolution tree construction showed that species A1 diverged before the last common ancestor of *V. alfalfae*, *V. dahliae* and *V. nonalfalfae*, whereas species D1 only recently diverged from *V. dahliae* as previously reported (Inderbitzin et al. 2011a). Intriguingly, the phylogenetic positions of the parental genomes did not correspond to that of the mitochondrial genome of *V. longisporum* (Figure 6). Mitochondrial DNA is subjected to laws different from the Mendelian principles of segregation and independent assortment as it is maternally inherited in most sexual eukaryotes, including numerous fungal species (Basse 2010). However, for particular organisms, bi-parental mitochondrial inheritance is common. For instance, in *S. cerevisiae* heteroplasmy can be maintained for ~20 generations, allowing parental mitochondrial genomes to recombine (Fritsch et al. 2014). The mitochondrial genome of *V. longisporum* has been inherited bi-parentally, as the mitochondrial genome is a mosaic consisting of alternating regions derived from the A1 or D1 parent (Figure 6). Similar to *V. longisporum*, two incipient species of the budding yeast *Saccharomyces paradoxus* contributed to the mitochondrial genome of their natural hybrid offspring (Leducq et al. 2017). Bi-parental inheritance is often associated with hybridization, although it is uncertain if hybridization facilitates paternal leakage or if bi-parental inheritance is a common phenomenon that is easer to detect in hybrids (Barr et al. 2005).

## Conclusion

The *V. longisporum* genome consists of two near to complete genomes of its haploid parents. Rearrangements between these parental chromosome sets occurred, resulting in a mosaic genome structure. *V. longisporum* genomes display high plasticity, as 60 synteny breaks were confirmed between strains with high nucleotide identity. Conceivably, the absence of meiotic constraints and the presence of orthologous DNA clusters provide ample opportunities for the *V. longisporum* genome to recombine. Inter-parental genomic recombination and functional diversification of homeologs give *V. longisporum* an additional potential to adapt to environmental alterations, which haploid *Verticillium* spp. do not have. This may have enabled *V. longisporum* to alter its host range and cause disease on Brassicaceous plants.

## ACKNOWLEDGEMENTS

The authors would like to thank the Marie Curie Actions programme of the European Commission that financially supported the research of J.R.L.D. Work in the laboratories of B.P.H.J.T. and M.F.S is supported by the Research Council Earth and Life Sciences (ALW) of the Netherlands Organization of Scientific Research (NWO).

## SUPPLEMENTARY MATERIAL

**Fig. S1:**
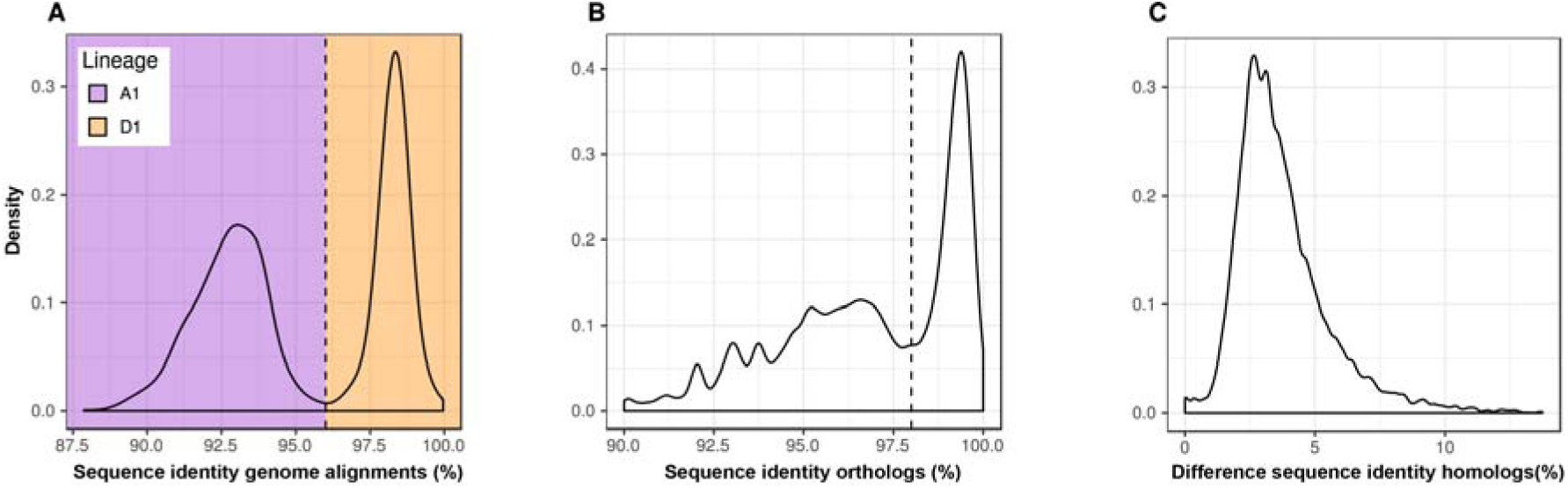
Lines of evidence for the parental origin of *V. longisporum* genomic regions. (A) Distribution of sequence identity of *V. longisporum* alignments to *V. dahliae*. (B) The distribution of the sequence identity between *V. longisporum* exonic regions of genes and their *V. dahliae* orthologs. (C) Distribution of sequence identity between exonic regions of *V. longisporum* homologs that are present in two copies. Strains VLB2 and JR2 were used for *V. longisporum* and *V. dahliae*, respectively.

**Fig. S2:**
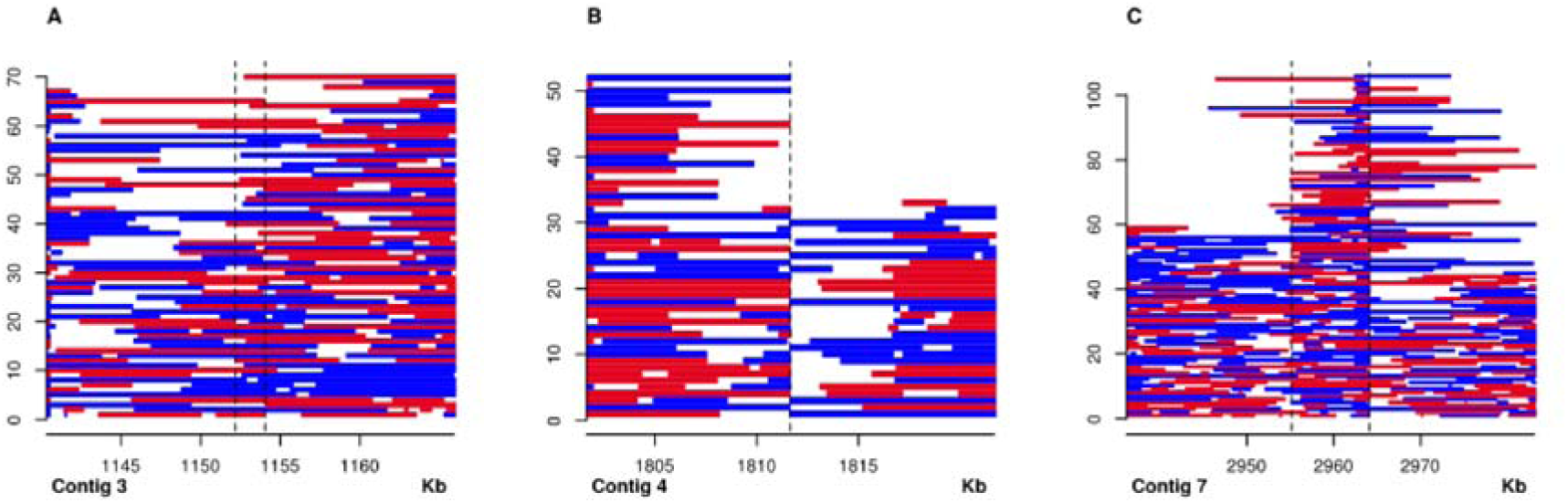
Synteny break confirmation by mapping *V. longisporum* VLB2 reads to the VL20 genome. Red and blue bars represent forward and reverse aligned VLB2 reads to theVL20 genome. The dashed lines show the suggested position in synteny break through genome alignment. (A) In particular cases, read alignments did not confirm the break in synteny. (B) In other cases, breaks in synteny were confirmed as reads abruptly stopped and started on these genome positions. (C) Breaks were also considered truthful if regions showed overlap in repeat-rich regions where read overlap between adjacent genome regions is lacking.

**Fig. S3:**
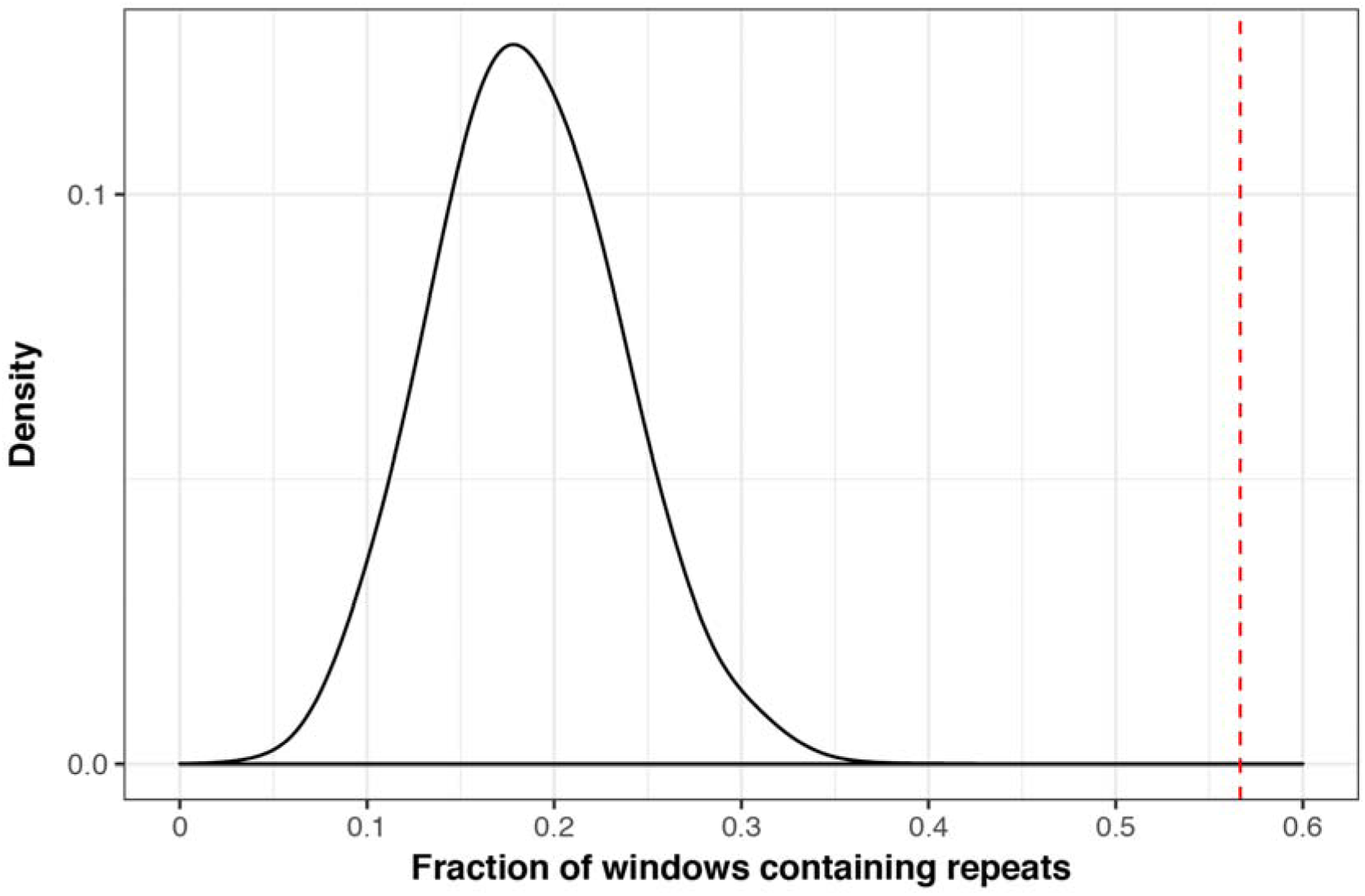
The association of synteny breaks with repetitive elements. The black curve represents the fraction of 60 randomly chosen 1 kb windows in the *V. longisporum* VL20 that are repeat-rich, which has been permutated 10,000 times. The red line indicates the fraction of true breaks that lay in a 1 kb window enriched for repeats (57%).

